# Genomic data suggest parallel dental vestigialization within the xenarthran radiation

**DOI:** 10.1101/2022.12.09.519446

**Authors:** Christopher A. Emerling, Gillian C. Gibb, Marie-Ka Tilak, Jonathan J. Hughes, Melanie Kuch, Ana T. Duggan, Hendrik N. Poinar, Michael W. Nachman, Frédéric Delsuc

## Abstract

The recent influx of genomic data has provided greater insights into the molecular basis for regressive evolution, or vestigialization, through gene loss and pseudogenization. As such, the analysis of gene degradation patterns has the potential to provide insights into the evolutionary history of regressed anatomical traits. We specifically applied these principles to the xenarthran radiation (anteaters, sloths, armadillos), which is characterized by taxa with a gradation in regressed dental phenotypes. Whether the pattern among extant xenarthrans is due to an ancient and gradual decay of dental morphology or occurred repeatedly in parallel is unknown. We tested these competing hypotheses by examining 11 core dental genes in most living species of Xenarthra, characterizing shared inactivating mutations and patterns of relaxed selection during their radiation. Here we report evidence of independent and distinct events of dental gene loss in the major xenarthran subclades. First, we found strong evidence of complete enamel loss in the common ancestor of sloths and anteaters, suggested by the inactivation of five enamel-associated genes (*AMELX, AMTN, MMP20, ENAM, ACP4*). Next, whereas dental regression appears to have halted in sloths, presumably a critical event that ultimately permitted adaptation to an herbivorous lifestyle, anteaters continued losing genes on the path towards complete tooth loss. Echoes of this event are recorded in the genomes of all living anteaters, being marked by a 2-bp deletion in a gene critical for dentinogenesis (*DSPP*) and a putative shared 1-bp insertion in a gene linked to tooth retention (*ODAPH*). By contrast, in the two major armadillo clades, genes pertaining to the dento-gingival junction and amelogenesis appear to have been independently inactivated prior to losing all or some enamel. These genomic data provide evidence for multiple pathways and rates of anatomical regression, and underscore the utility of using pseudogenes to reconstruct evolutionary history when fossils are sparse.

## Introduction

Regressive evolution involves the vestigialization or loss of formerly adaptive traits over time, with examples ranging from the reduction of wings in flightless birds, the degeneration of limbs in various squamates and aquatic mammals, and the degradation of eyes in species occupying extreme dim-light niches. The reduction of such traits is often thought to result from a lack of adaptive utility after entering a novel niche, resulting in relaxed selection and regression through drift, and/or direct selection against the trait to minimize energetic costs (Fong et al., 1995; Jeffery, 2009; Lahti et al., 2009). The increasing availability of genomic data has permitted the documentation of parallel, and possibly causal, mutational events in genes associated with these and other vestigial phenotypes, providing a greater understanding of the genetic basis of regressive evolution (Albalat and Cañestro, 2016; Burga et al., 2017; Emerling, 2017; Emerling and Springer, 2014; Leal and Cohn, 2016; Sharma et al., 2018).

For instance, while the teeth of gnathostomes can be considered a key adaptive trait in the origin of and radiation of this clade, dentition has subsequently regressed in numerous lineages (Charles et al., 2013; Davit-Béal et al., 2009), such as baleen whales, turtles, birds and pangolins. Notably, there are well-documented corresponding genomic signals behind these numerous instances of dental degeneration. Genetic association studies in humans with dental diseases and mouse knockout models have led to a robust understanding of the genetics underlying tooth development (Meredith et al., 2014; Smith et al., 2017), and comparative genomic analyses of edentulous (toothless) and enamelless vertebrates have revealed that many of these same dental genes were deleted or have eroded into unitary pseudogenes. Indeed, the list of documented dental pseudogenes in such vertebrates includes those encoding (1) enamel matrix proteins (EMPs), which provide a protein scaffold for the seeding of hydroxyapatite crystals during enamel development (ENAM [enamelin], AMELX [amelogenin], AMBN [ameloblastin]) (Choo et al., 2016; Delsuc et al., 2015; Meredith et al., 2014, 2013, 2009; Sire et al., 2008), (2) a metalloproteinase that processes these matrix proteins into their mature forms (MMP20 [enamelysin]) (Meredith et al., 2011a, 2014), (3) other proteins expressed in both enamel-forming ameloblasts and enamel-contacting gingiva (AMTN [amelotin], ODAM [odontogenic ameloblast-associated]) (Gasse et al., 2012; Meredith et al., 2014; Springer et al., 2019), (4) proteins of unknown function but showing clear associations with enamel formation (ACP4 [acid phosphatase 4; formerly called ACPT], ODAPH [odontogenesis-associated phosphoprotein; formerly called *C4orf26*]) (Mu et al. 2021; Sharma et al., 2018; Springer et al., 2016), and (5) a protein that contributes to the dentin matrix (DSPP [dentin sialophosphoprotein]) (Meredith et al., 2014; Sire et al., 2008; McKnight and Fisher 2009).

Given the parallel signals between genes and anatomy during the evolution of degenerated traits, genomic data have the potential to provide insights into the sequence, dynamics and consequences of regressive evolution, particularly in cases where the fossil record is limited. Xenarthra, which includes armadillos (Cingulata), sloths (Folivora) and anteaters (Vermilingua), represents a compelling example of the process of regressive evolution in regards to dentition. In contrast to the completely edentulous clades of baleen whales, birds, turtles and pangolins, xenarthrans span a spectrum of regressed dental phenotypes across a continual 68 million year radiation (Charles et al., 2013; Ciancio et al., 2014; Davit-Béal et al., 2009; Gibb et al., 2016; Vizcaíno, 2009). For instance, anteaters are united with sloths in the clade Pilosa, but while anteaters entirely lack teeth, sloths have intermediately-regressed teeth, possessing an edentulous premaxilla and simple, peg-like, single-rooted, enamelless dentition. Indeed, their teeth have deviated so far from the ancestral tribosphenic form, that only recent developmental research has provided evidence of dental homologies to other placental mammals (Hautier et al., 2016). Sister to Pilosa are the Cingulata, which are divided into the extant families Dasypodidae and Chlamyphoridae. Most extant armadillos have an edentulous premaxilla, as well as peg-like, single-rooted teeth, which lack enamel in adult animals (Ferigolo, 1985). A notable exception is Dasypodidae, which possess vestigial enamel on deciduous teeth and prismatic or prismless enamel on permanent teeth of juveniles, which wears away with use (Ciancio et al., 2021; Martin, 1916; Spurgin, 1904).

The spectrum of degenerated dental phenotypes across Xenarthra raises a number of distinct questions. The first set concerns the specific history of the xenarthran clade. Anteaters are myrmecophagous mammals, almost exclusively consuming copious amounts of ants and termites by employing an extensive tongue with sticky saliva, and as such have little use for teeth. Sloths, by contrast, are herbivorous and often folivorous, and need to chew fibrous material with their seemingly ill-equipped, enamelless teeth. Among armadillos, some are relatively myrmecophagous (e.g., tolypeutines), while others are more omnivorous (e.g., euphractines). Whereas an extended history of myrmecophagy is universally associated with dental regression (Charles et al., 2013; Davit-Béal et al., 2009; Reiss, 2001), herbivory and omnivory are not. Accordingly, is the dental regression seen in xenarthrans the result of inheriting regressed teeth from their last common ancestor, which possibly had an insectivorous/myrmecophagous diet followed by subsequent dietary shifts to herbivory and omnivory? Or does it represent parallel events in multiple lineages?

A second set of questions concerns the timing and patterns of regressive evolution. First, has the degeneration of teeth in xenarthrans taken place over a short period of time, consistent with selection against their presence, or has it been a gradual process over many millions of years in a manner more in line with relaxed selection and genetic drift? Furthermore, is there any sort of consistency in the sets of genes that are lost and the timing of those losses, or does the regression of teeth occur via divergent genomic patterns?

To answer these evolutionary questions, we collected genomic data to study patterns of pseudogenization and selection pressure in 11 core dental genes for most living species of Xenarthra. Our results point to xenarthran teeth having repeatedly regressed in parallel, showing distinct patterns of gene loss in different lineages in order to give rise to the variation in dentition observed across the clade today. They further suggest that regressive evolution can take place both gradually and in relatively rapid, discrete phases for the same trait during the radiation of a single clade.

## Methods

To study patterns of dental gene loss in xenarthrans, we assembled a dataset of 11 dental genes: nine genes have well-characterized functions and/or expression patterns tied to tooth development and are frequently pseudogenized in edentulous and enamelless taxa (*ACP4, AMBN, AMELX, AMTN, DSPP, ENAM, MMP20, ODAM, ODAPH*; Choo et al., 2016; Delsuc et al., 2015; Gasse et al., 2012; McKnight and Fisher, 2009; Meredith et al., 2014, 2013, 2011a, 2009; Mu et al., 2021; Sharma et al., 2018; Sire et al., 2008; Smith et al., 2017; Springer et al., 2019, 2016), and two other genes (*DMP1, MEPE*) are expressed in dentin (Sun et al., 2011; Gullard et al., 2016).. Our taxonomic coverage included 31 xenarthran species (four anteaters, six sloths, seven dasypodid armadillos, 14 chlamyphorid armadillos) plus 25 outgroup species spanning the remaining three superorders of placental mammals. We used a combination of strategies to reconstruct gene sequences: targeted sequencing of PCR amplified regions, exon-capture, whole-genome sequencing, and retrieval of sequences from publicly available genome assemblies (Supplementary Tables S1, S2, Figure S1).

### Biological samples

Xenarthran tissue samples used for DNA extractions and amplifications of dental gene exons came from the Animal Tissue Collection of the Institut des Sciences de Montpellier (Supplementary Table S1): nine-banded armadillo (*Dasypus novemcinctus* ISEM T-JL556), greater long-nosed armadillo (*Dasypus kappleri* ISEM T-2977), southern naked-tailed armadillo (*Cabassous unicinctus* ISEM T-2291), large hairy armadillo (*Chaetophractus villosus* ISEM NP390), giant armadillo (*Priodontes maximus* ISEM T-2353), southern three-banded armadillo (*Tolypeutes matacus* ISEM T-2348), pichi (*Zaedyus pichiy* ISEM T-6060), pink fairy armadillo (*Chlamyphorus truncatus* ISEM T-CT1), pale-throated three-fingered sloth (*Bradypus tridactylus* ISEM T-1476), Linneaus’s two-fingered sloth (*Choloepus didactylus* ISEM T-1722), Hoffmann’s two-fingered sloth (*Choloepus hoffmanni* ISEM T-6052), southern tamandua (*Tamandua tetradactyla* ISEM T-6054), giant anteater (*Myrmecophaga tridactyla* ISEM T-2862), and pygmy anteater (*Cyclopes didactylus* ISEM T-1631), and from the Museum of Vertebrate Zoology (Berkeley, CA, USA) for the brown-throated three-fingered sloth (*Bradypus variegatus* MVZ 155186). Xenarthran specimens and corresponding Illumina genomic libraries used in exon capture experiments were those previously generated in Gibb et al. (2016). The greater fairy armadillo museum sample (*Calyptophractus retusus* ZSM T-Bret) and the pink fairy armadillo (*Chlamyphorus truncatus* ISEM T-CT1) used for whole genome shotgun sequencing were respectively obtained from the Bavarian State Collection of Zoology (Munich, Germany) and provided by Dr. Mariella Superina as previously detailed in Delsuc et al. (2012). The pygmy anteater (*Cyclopes didactylus* JAG M2300), the pale-throated three-fingered sloth (*Bradypus tridactylus* JAG M1664), the giant armadillo (*Priodontes maximus* M844), and the greater long-nosed armadillo (Dasypus kappleri M3462) used for whole genome shotgun sequencing came from the JAGUARS animal tissue collection hosted at the Institut Pasteur de la Guyane (Cayenne, French Guiana). The six-banded armadillo (*Euphractus sexcinctus* T-ESE1) used for whole genome shotgun sequencing was sampled at the Zoo de Lunaret (Montpellier, France). Finally, the southern naked-tailed armadillo (*Cabassous unicinctus* MVZ 155190) sample used for whole genome assembly was derived from a frozen tissue sample from the Museum of Vertebrate Zoology (Berkeley, CA, USA). In accordance with the policy of sharing benefits and advantages (APA; TREL1916196S/224), biological material from French Guiana collected after October 2014 has been registered in the JAGUARS collection supported by Kwata NGO, Institut Pasteur de la Guyane, DEAL Guyane, and Collectivité Territoriale de la Guyane. Biological samples from the JAGUARS collection were exchanged through formal material transfer agreements granted by DEAL Guyane.

### DNA extractions, PCR amplifications, and Sanger sequencing

Total genomic DNA was extracted from tissue samples (Supplementary Table S1) preserved in 95% ethanol using the QIAampDNA extraction kit (Qiagen). Polymerase chain reaction (PCR) was used to target and amplify *AMBN, AMELX, DMP1, DSPP, ENAM*, and *MEPE* exons. To do so, a number of primer pairs were designed from alignments of available sequences (Supplementary Table S3). PCR conditions were as follows: 95°C for 5 min, followed by 40 cycles at 95°C for 30 s, 50-55°C for 30 s, 72°C for 45 s, and a final extension step at 72°C for 10 min. All PCR products were then purified from 1% agarose gels using Amicon Ultrafree-DA columns (Millipore Corporation, Bedford, MA, USA) and sequenced on both strands using the polymerase chain reaction primers with the Big Dye Terminator cycle sequencing kit on an Applied ABI Prism 3130XL automated sequencer. Electropherograms were checked by eye and assembled into contigs using Geneious Prime (Kearse et al., 2012).

### Target sequence capture and sequencing

Baits for DNA sequence exon capture were designed using available xenarthran sequences for the complete CDSs of all focal dental genes except *ACP4*. This was due to its link to amelogenesis imperfecta being reported after the design and synthesis of these baits. For each gene, 80mer baits were generated with a 4x tiling density and were then BLASTed against the genome assemblies of *Choloepus hoffmanni* (GCA_000164785.2) and *Dasypus novemcinctus* (GCA_000208655.2). Baits with more than one hit and a Tm outside the range 35-40°C were excluded. This resulted in a final set of 5,262 baits (Supplementary Dataset S1) that were synthesized as part of a myBaits RNA kit by Arbor Biosciences (Ann Arbor, MI, USA). The xenarthran Illumina libraries previously prepared (Gibb et al., 2016) were subsequently enriched with the designed bait set in order to capture target sequences following previously described methodological procedures (Delsuc et al., 2018). All enriched libraries were pooled together at varying concentrations with the aim of producing one million reads for sequencing. Sequencing of the enrichment set was performed at McMaster Genomics Facility (McMaster University, Hamilton, ON, Canada) on an Illumina MiSeq instrument using 150 bp paired-end reads.

Index and adapter sequences were removed from raw reads and overlapping pairs merged with leeHom (Renaud et al., 2014), and then mapped to all xenarthran reference sequences available with a modified version of BWA (Li and Durbin, 2009; Stenzel, 2017). Mapped reads were additionally filtered to those that were either merged or properly paired, had unique 5’ and 3’ mapping coordinates, and then restricted to reads of at least 24 bp with SAMtools (Li et al., 2009). The bam files were then imported into Geneious to select the best assembly for each sequence depending on the most successful reference sequence. All sequences have been deposited in GenBank at the National Center for Biotechnology Information (NCBI) under accession numbers OP966064 to OP966335.

### Whole genome shotgun libraries construction and sequencing

For the greater fairy armadillo museum specimen (*Calyptophractus retusus* ZMS T-Bret), we constructed new whole genome DNA Illumina libraries using a previously extracted genomic DNA and library preparation procedure (Gibb et al., 2016). These degraded DNA libraries were sequenced at the Vincent J. Coates Genomics Sequencing Laboratory (University of California, Berkeley, CA, USA) on an Illumina HiSeq4000 instrument using 50 bp single reads. Whole genome DNA Illumina library preparation and sequencing of *Bradypus tridactylus* (JAG M1664), *Cyclopes didactylus* (JAG M2300), *Chlamyphorus truncatus* (ISEM T-CT1), *Euphractus sexcinctus* (ISEM T-ESE1), *Dasypus kappleri* (JAG M3462), and *Priodontes maximus* (JAG M844) were outsourced to Novogene Europe (Cambridge UK). These libraries were sequenced on an Illumina NovaSeq instrument using 150 bp paired-end reads. The resulting short reads were cleaned from sequencing indexes and adaptors and quality filtered using Trimmomatic (Bolger et al., 2014) with default parameters. These were then mapped to their closest available relative xenarthran reference sequence using the Geneious mapping algorithm with default settings. Consensus sequences of mapped reads were called using the 50% majority rule for each targeted gene.

### Whole genome sequencing and assemblies

For the southern naked-tailed armadillo (*Cabassous unicinctus* MVZ 155190), a PCR-free library Illumina library with 450 bp inserts was prepared according to the recommendations of the Broad Institute for subsequent genome assembly with DISCOVAR de novo (https://software.broadinstitute.org/software/discovar/blog/?page_id=375) following the strategy of the Zoonomia project (Zoonomia consortium, 2020). The library was sequenced at the Vincent J. Coates Genomics Sequencing Laboratory (University of California, Berkeley, CA, USA) on an Illumina HiSeq4000 instrument using 250 bp paired-end reads. A draft genome assembly was produced from these data using the DISCOVAR de novo assembler (https://github.com/broadinstitute/discovar_de_novo/releases/tag/v52488) using the following command: DiscovarDeNovo READS=[ReadsFile] OUT_DIR=[OutputDirectory] NUM_THREADS=24 MAX_MEM_GB=700.

We also utilized multiple whole genome assemblies deposited in NCBI Whole Genome Shotgun (WGS) contig database (Supplementary Table S2). To obtain genes on NCBI, we BLASTed human reference nucleotide sequences, composed of exons, introns and flanking sequences, against the WGS database using discontiguous megablast (BLAST+ 2.8.0). We then used the NCBI-derived xenarthran sequences to BLAST using discontiguous megablast against Discovar de novo assemblies imported into Geneious. Sequences derived from assemblies that contained strings of more than 10 unknown nucleotides (Ns) had these stretches of Ns reduced to 10 for ease of alignment. Raw sequencing reads have been deposited in the Short Read Archive (SRA) at the National Center for Biotechnology Information (NCBI) under Bioproject PRJNA907496. The Discovar de novo draft genome assembly for the southern naked-tailed armadillo is available on NCBI (JAQZBX000000000).

### Dataset assembly

Xenarthran sequences were assembled, and occasionally combined, from the various methodologies employed (Supplementary Datasets S2–12). Outgroup sequences were retrieved by BLASTing reference sequences against GenBank using discontiguous megablast and obtaining NCBI annotated gene models (Supplementary Table S2). All orthologs of xenarthrans and outgroup sequences were aligned using MUSCLE (Edgar, 2004) in Geneious, examined by eye and adjusted manually to correct for errors in the automated alignments. After characterizing putative inactivating mutations (see below), we prepared codon alignments suitable for subsequent selection pressure analyses based on dN/dS estimations (Supplementary Datasets S13–23) by removing insertions, incomplete codons and stop codons from individual sequences, and any other portions of sequences with dubious homology, including: exons 7–9 of *AMBN*, given probable exon duplications within the human reference (Toyosawa et al., 2000); *DSPP* highly repetitive region at the 3’ end of exon 4 (McKnight and Fisher 2009; Fisher, 2011); and exon 2 of *ODAPH* in humans, which is a splice variant that is not common to all placental mammals (Springer et al., 2016). The evolutionary history of each gene was estimated by maximum likelihood phylogenetic reconstruction using RAxML v8.2.11 (Stamatakis, 2014) within Geneious (GTR+GAMMA model, Rapid hill-climbing) to detect aberrant sequences (e.g., possible contaminants and annotation errors stemming from poor gene models).

### Inactivating mutations and dN/dS ratio analyses

We classified dental genes in five functional categories: (1) enamel matrix (*AMELX, AMBN, ENAM*), (2) enamel matrix-processing (*MMP20*), (3) dento-gingival junction (*AMTN, ODAM*), (4) unknown enamel function (*ACP4, ODAPH*) and (5) dentinogenesis (*DSPP, DMP1, MEPE*). We searched for putative inactivating mutations, including frameshift insertions or deletions, premature stop codons, start codon mutations, and splice site mutations, and noted any mutations shared by multiple species within a clade. We also noted examples in which exons were not recovered, some of which may correspond to whole exon deletions, although validation would require more contiguous genome assemblies. We then performed several analyses to estimate the pattern and timing of shifts in selection pressure in tooth genes among our focal xenarthran taxa.

We first reconstructed the evolution of the dN/dS ratio (ω) for each gene using the Bayesian approach implemented in Coevol 1.4b (Lartillot and Poujol, 2011). Coevol provides a visual representation of the variation in dN/dS ratio estimates across a phylogeny (e.g. Lartillot and Delsuc, 2012; Springer et al., 2019). We used the *dsom* procedure to jointly estimate branch specific dN/dS ratios, divergence times, body sizes, generation times, and ages at sexual maturity modeled as a multivariate Brownian diffusion process across the phylogeny. We used the same composite placental mammal topology for all analyses (Emerling et al., 2015; Gibb et al., 2016), assumed fossil calibrations from previous studies (Meredith et al., 2011b; Emerling et al, 2015; Foley et al., 2016), and extracted the three life history traits (body size, generation time, and age at sexual maturity) from the PanTHERIA database (Jones et al., 2009). We set the prior on the root node to 97 Ma with a standard deviation of 20 Ma following the molecular dating estimates of Meredith et al. (2011b). For each dataset, we ran two independent MCMC for a total of 1000 cycles, sampling parameters every cycle. MCMC convergence was checked by monitoring the effective sample size of the different parameters using the *tracecomp* command of Coevol. The first 100 points of each MCMC were excluded as burnin and posterior inferences were made from the remaining 900 sampled points of each chain.

Next, based on the patterns of inactivating mutations, we performed branch model dN/dS ratio analyses (Yang 1998; Yang and Nielsen, 1998) using *codeml* in PAML v4.8 (Yang, 2007) to estimate the selection pressure experienced by the different tooth genes throughout xenarthran history. We used the same topology as for Coevol estimations in all analyses and employed the following protocol for each gene, which is summarized graphically in Supplementary Figure S2. First, we ran *codeml* with a one-ratio model, successively employing codon frequency models 0, 1 and 2, then using the Akaike information criterion to determine the best-fitting model (Supplementary Table S4). Next, using the branch model approach, we allowed ω to vary across the phylogeny in a multi-ratio model with branch labels set for (a) each set of branches within a clade that post-date the minimum gene inactivation date inferred from shared inactivation mutations, (b) each branch that coincides with inferred gene inactivation, and (c) certain branches that predate gene inactivation, often grouped together with multiple such branches. Note that this does not correspond to a free ratio model, but rather follows a previously described approach (Meredith et al., 2009). In every case, when taxon representation was sufficient, we set separate branch categories for stem Xenarthra, stem Pilosa, stem Cingulata, stem Chlamyphoridae, and stem Dasypodidae. All non-xenarthran branches were grouped together in a single ω category. Given that all selected outgroup taxa have teeth with enamel, this background dN/dS ratio was assumed to be the average baseline (background) ω estimate for functional dental genes. Finally, subsequent models were run in which focal xenarthran branches were fixed as part of this background ratio and then 1, respectively, to test if they diverged significantly from these contrasting assumptions of purifying selection (background) versus relaxed selection (ω = 1). As an example, the stem xenarthran branch was freely estimated in the initial multi-ratio model, then a second model fixed the stem xenarthran branch as part of the background ratio, and then a third model fixed the stem xenarthran branch at 1. This was carried out for all non-background branches. Models were then compared statistically using likelihood-ratio tests (Yang, 2007).

## Results

### Widespread pseudogenisation of dental genes in xenarthrans

Nine out of the 11 examined tooth genes are inactivated in most xenarthrans, though there are different patterns distinct to certain clades (Figure 1; Supplementary Tables S5-S15). The toothless anteaters have the greatest proportion of pseudogenized genes (82%), followed by the enamelless sloths (55%-64%) and chlamyphorid armadillos (45-55%), and the weakly-enamelled dasypodid armadillos (27-45%).

**Figure 1:**
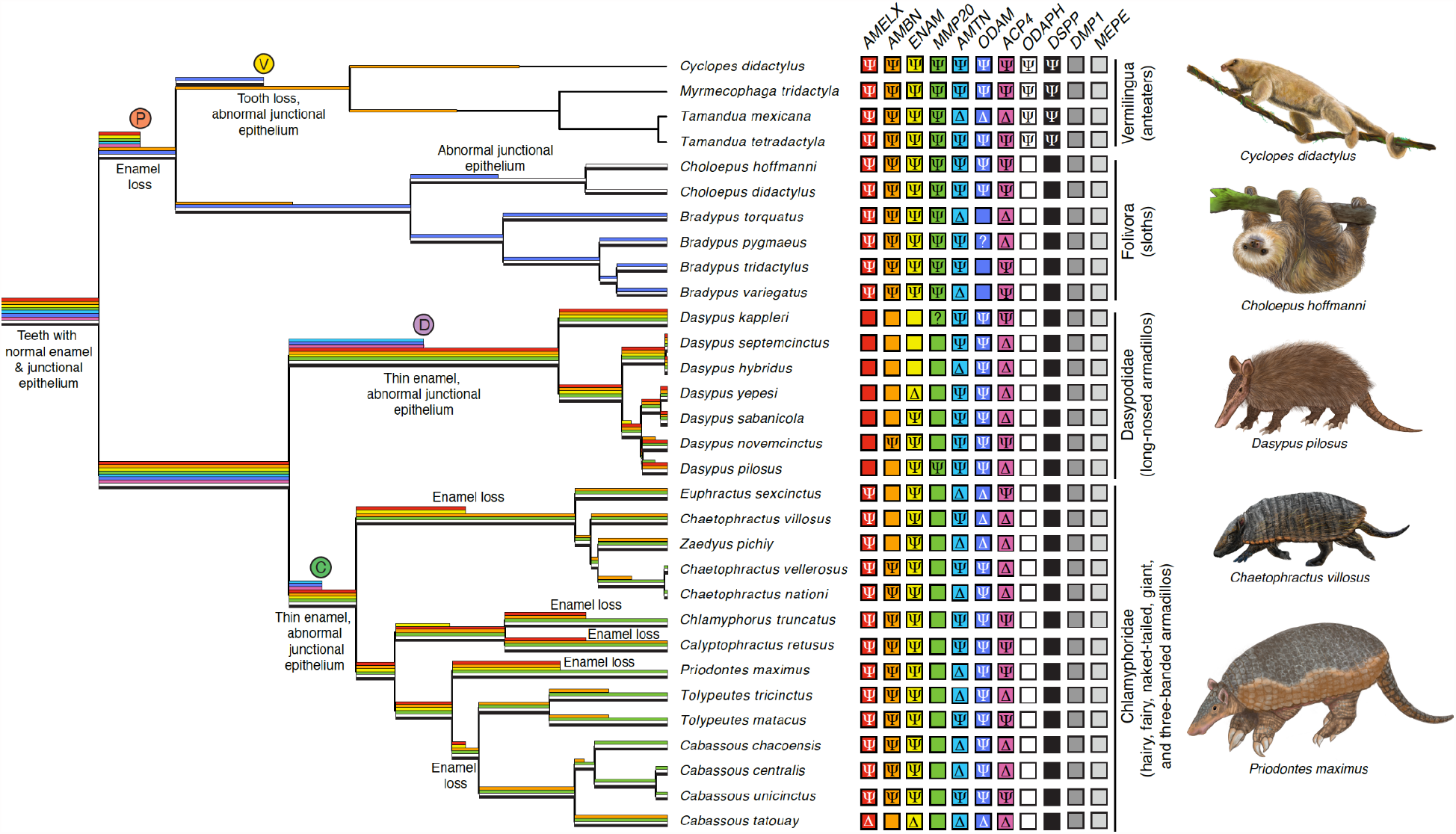
Dental gene inactivations across Xenarthra indicating distribution of gene losses (right) and inferred regressive dental events based on shared inactivating mutations (SIMs) of key genes (left). Thin enamel = evidence of inactivation in *ACP4*; abnormal junctional epithelium = evidence of inactivation in both *AMTN* and *ODAM*; enamelless teeth = inactivation of *AMELX*; edentulous = inactivation of *ODAPH* and *DSPP* (see text for interpretations). Branches on which gene inactivations occurred are inferred from SIMs, which are summarized in the Results, Figures 2 and 3 and Supplementary Tables S5–S15. Gene losses in right columns are indicated by the following: Ψ = positive evidence of pseudogenization; Δ = no positive evidence of pseudogenization, but gene inactivation inferred from phylogenetic distribution of shared mutations; ? = no data; empty box = gene putatively intact. Timing of gene inactivations arbitrarily placed midway on the corresponding branch, indicated by termination of colored branch. Colors on branches correspond to genes using the same colors on the right (e.g., *AMELX* = red, *AMBN* = orange, etc.). Colored circles with letters P, V, C and D correspond with letters in Figures 2 and 3. Paintings by Michelle S. Fabros.

Genes encoding the enamel matrix proteins (EMPs) are commonly inactivated. Among all pilosans (anteaters and sloths), *AMBN, AMELX* and *ENAM* are pseudogenes (Figure 2). Among chlamyphorids, *ENAM* and *AMELX* are inactivated in all species, whereas *AMBN* varies, appearing intact in some euphractines. The sequences for the latter remain incomplete, however, raising the possibility of undetected inactivating mutations. Within dasypodids, EMPs are largely intact: *ENAM* is variable in its inactivation, *AMELX* is never a pseudogene and *AMBN* inactivation is only suggested by a start codon mutation in the nine-banded armadillo (*Dasypus novemcinctus*).

**Figure 2:**
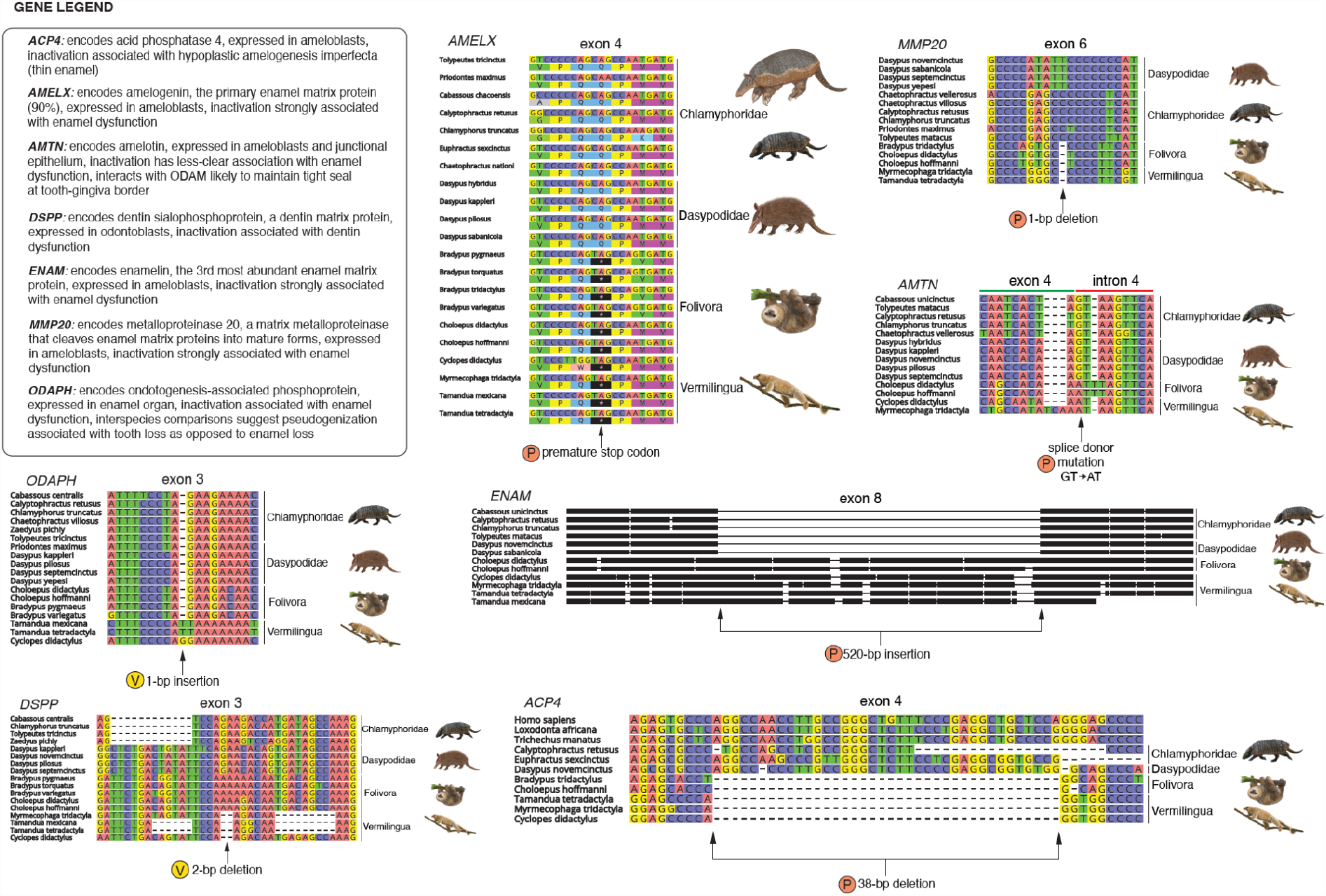
DNA alignments providing examples of shared inactivating mutations indicating key dental gene losses in pilosans. Colored circles with the letters P and Vcorrespond with letters in Figure 1. P = gene inactivation inferred on stem Pilosa branch; V = gene inactivation inferred on stem Vermilingua branch. Paintings by Michelle S. Fabros.

The metalloproteinase-encoding *MMP20* gene is clearly pseudogenized in all pilosan species (Figure 2). Among armadillos, the data suggest a very different trend: only two species provide evidence of inactivation, with the hairy long-nosed armadillo (*Dasypus pilosus*) having a single stop codon in exon 2, and the northern long-nosed armadillo (*D. sabanicola*) possessing a polymorphic stop codon in exon 8. Missing data is an issue for some exon capture sequences, but complete sequences are provided for species with genome assemblies, suggesting that *MMP20* is indeed intact in many if not the vast majority of armadillos, a conclusion further supported by selection pressure analyses based on dN/dS ratio (see below).

The genes encoding the ameloblast and gingiva-expressed amelotin (*AMTN*) and odontogenic ameloblast-associated protein (*ODAM*) are almost universally inactivated in xenarthrans (Figure 1–3). The lone exception is *ODAM* in the three-fingered sloths (*Bradypus* spp.), all of which fail to show evidence of inactivation.

Among the two genes encoding proteins with less clear roles in enamel formation (*ACP4, ODAPH*), only *ACP4* shows evidence of widespread pseudogenization, being unambiguously inactivated in every species for which we had sufficient data (Figures 2 and 3), and further supported by patterns of shared mutations (see below). By contrast, *ODAPH* is only inactivated in anteaters (Figure 2).

**Figure 3:**
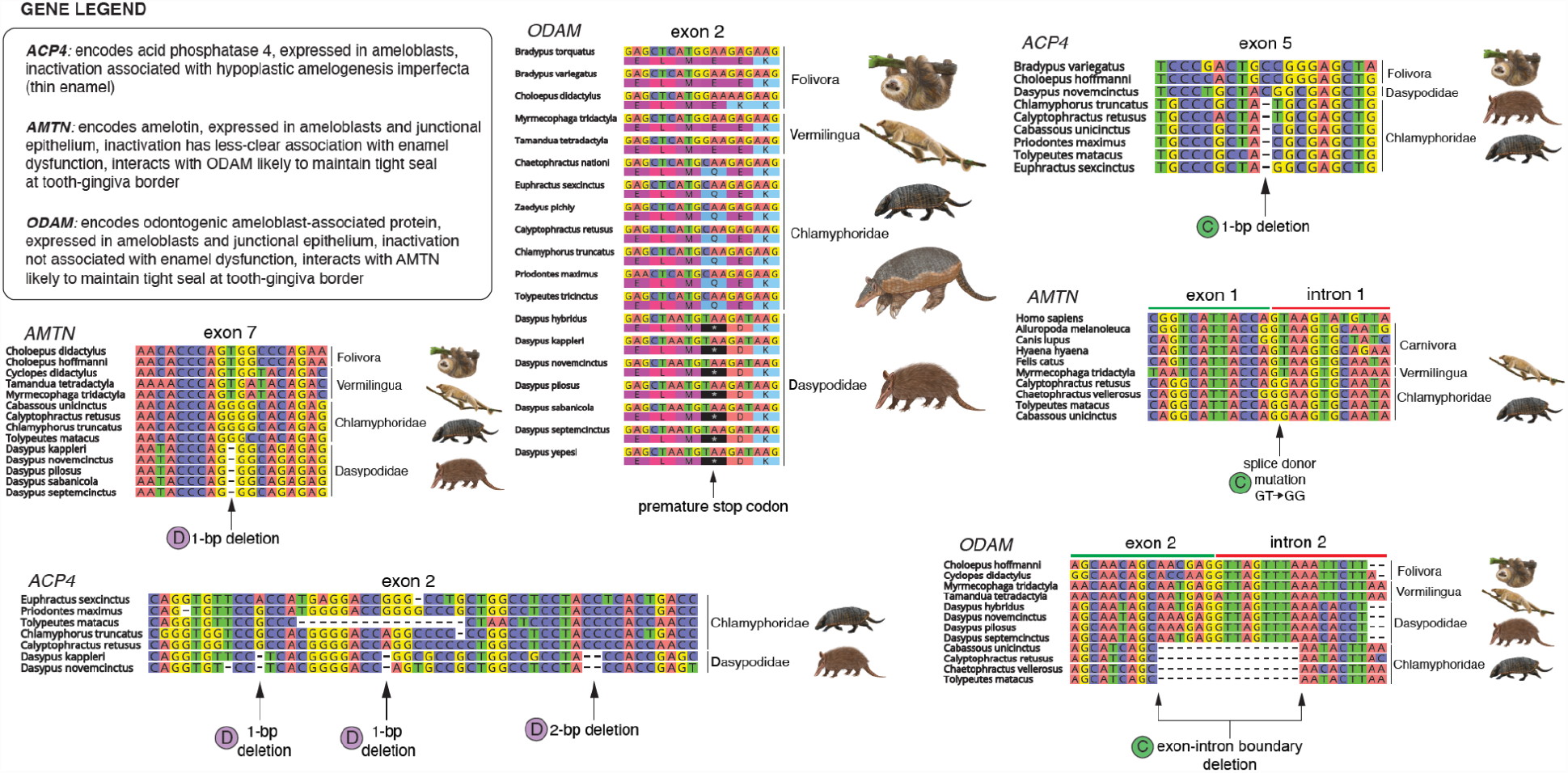
DNA alignments providing examples of shared inactivating mutations indicating key dental gene losses in armadillos. Colored circles with letters C and D correspond with letters in Figure 1. C = gene inactivation inferred on stem Chlamyphoridae branch; D = gene inactivation inferred on stem Dasypodidae branch. Paintings by Michelle S. Fabros.

Finally, among the genes encoding dentinogenesis proteins (*DSPP, DMP1, MEPE*), only *DSPP* shows evidence of inactivation, and, again, solely among the anteaters (Figure 2). *DMP1* and *MEPE* have important roles in bone formation (Gullard et al., 2016; Sun et al., 2011), which may explain their consistent retention.

### Reconstructing patterns of shared inactivating mutations

The presence of shared inactivating mutations (SIMs) provides strong evidence for the minimum date of pseudogenization in a lineage. If, for example, a frameshift indel or premature stop codon occurs at the same position in all species within a clade, then it is more parsimonious to assume that it was inherited from a common ancestor rather than being independently derived. Furthermore, multiple SIMs within a gene strengthen the case for shared history.

We found no unambiguous inactivating mutations shared among all xenarthran subclades. While we were unable to recover the complete coding sequence for every gene in every species, the phylogenetic distribution of taxa derived from whole genome sequencing means that this inference is unlikely to be the result of missing data (Supplementary Figures S3-S13). By contrast, we found multiple examples of SIMs among pilosans, dasypodids and chlamyphorids, respectively (Figures 2 and 3, Supplementary Tables S5, S7, S8, S11, S13, S14).

For pilosan-specific SIMS, all ten species we examined shared a premature stop codon in exon 4 of *AMELX*. Exon 4 of *AMTN* is followed by a splice donor mutation (GT → AT) shared by two-fingered sloths (*Choloepus* spp.) and the pygmy (*Cyclopes didactylus*) and giant (*Myrmecophaga tridactyla*) anteaters. Exon 8 of *ENAM* has three SIMs across pilosans, including a 1-bp deletion, premature stop codon and a roughly 520 bp insertion. Among the three sloths and two anteaters for which we assembled all of exon 6 of *MMP20*, they share a 1-bp deletion. For *ACP4*, we found evidence of a 38-bp deletion in exon 4 and two 1-bp deletions in exon 7 across four sloths and three anteaters (note: the latter alignment is ambiguous and may reflect a single 2-bp deletion). Finally, exon 3 of *AMBN* is preceded by a splice acceptor mutation (AG → AT) in sloths, but this exon is missing in anteaters. While this leaves open the possibility that this gene was inactivated in a stem pilosan, the dN/dS ratio results render this possibility unlikely (see below).

Two genes appear to have been inactivated at minimum in a stem or crown sloth lineage (Figure 1). All sloths possess three SIMs in *AMBN*, and two-fingered sloths possess a single SIM in *ODAM* (exon 8, 10-bp deletion), with three-fingered sloths lacking any discernible inactivating mutations in *ODAM*. Four genes appear to have been lost in the stem or a crown anteater lineage (Figure 1). Anteater SIMs include a stop codon in exon 9 of *ODAM*, a 1-bp insertion in exon 3 of *ODAPH*, and a 2-bp deletion in exon 3 of *DSPP* (Figure 2). By contrast, *AMBN* only clearly possesses SIMs (four) between the giant (*Myrmecophaga tridactyla*) and lesser anteaters (*Tamandua* spp.).

We found no evidence of SIMs shared among all armadillos, but *AMTN, ODAM* and *ACP4* all contain SIMs unique to each subclade (Figure 3). Among chlamyphorids, a chlamyphorine, euphractine and two tolypeutine armadillos possess a splice donor mutation in intron 1 of *AMTN* (GT → GG), and the same species share a 15-bp deletion of the splice donor region of intron 2 in *ODAM*. For *ACP4*, there is a 1-bp deletion in exon 5 and a splice acceptor mutation (GT → GG) in intron 5 among both chlamyphorines, three tolypeutines, and euphractines. Among dasypodids, a 1-bp deletion is found in exon 7 of *AMTN, ODAM* has premature stop codons in exons 4 and 9, and *ACP4* has at minimum 13 SIMS across at least five exons. Note that we did not recover exon 1 of *AMTN* in any dasypodids, which means that the splice donor mutation found in chlamyphorids could be shared among all armadillos. dN/dS analyses suggest this is a viable but uncertain possibility (see below).

Among the other genes, the pattern of SIMs in armadillos is much patchier (Figure 1). *AMBN*, for instance, only has shared mutations among congeners. By contrast, *AMELX* has SIMs in euphractines and tolypeutines, respectively, but none in chlamyphorines. Finally, *ENAM* possesses SIMs unique to all euphractines, chlamyphorines, and probably tolypeutines, respectively, and within dasypodids, *Dasypus novemcinctus* and *D. pilosus* share a splice donor mutation (AG → AT, intron 4) and *D. novemcinctus* and *D. sabanicola* share a 1-bp insertion (exon 8).

### Selection pressure analyses

Given that shared inactivating mutations provide only a minimum probable date for inactivation, they may underestimate the timing of the onset of relaxed selection on a gene. Gene dysfunction may predate the fixation of a frameshift indel or premature stop codon, e.g., due to disruption of non-coding elements. In order to evaluate changes in selection pressure resulting from dental regression, we first reconstructed the global variation in dN/dS across the placental phylogeny (Figure 4) using the program Coevol, which implements a form of rate smoothing through the incorporation of a Brownian motion model of continuous trait evolution. These analyses allow for the visualization and localization of shifts in selection pressure that have occurred within xenarthrans, likely corresponding to relaxed selection and pseudogenization events in the 11 focal dental genes.

**Figure 4:**
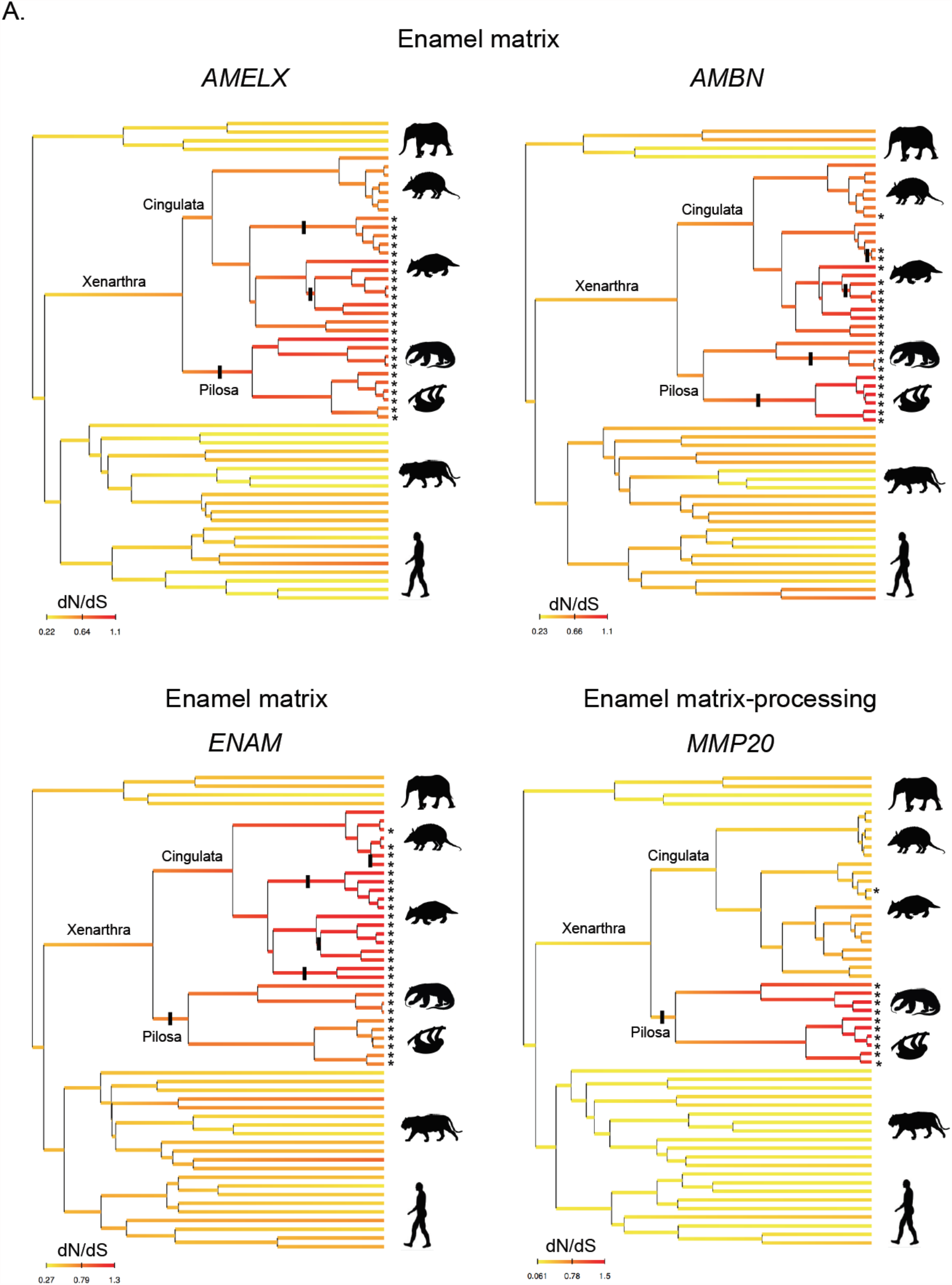
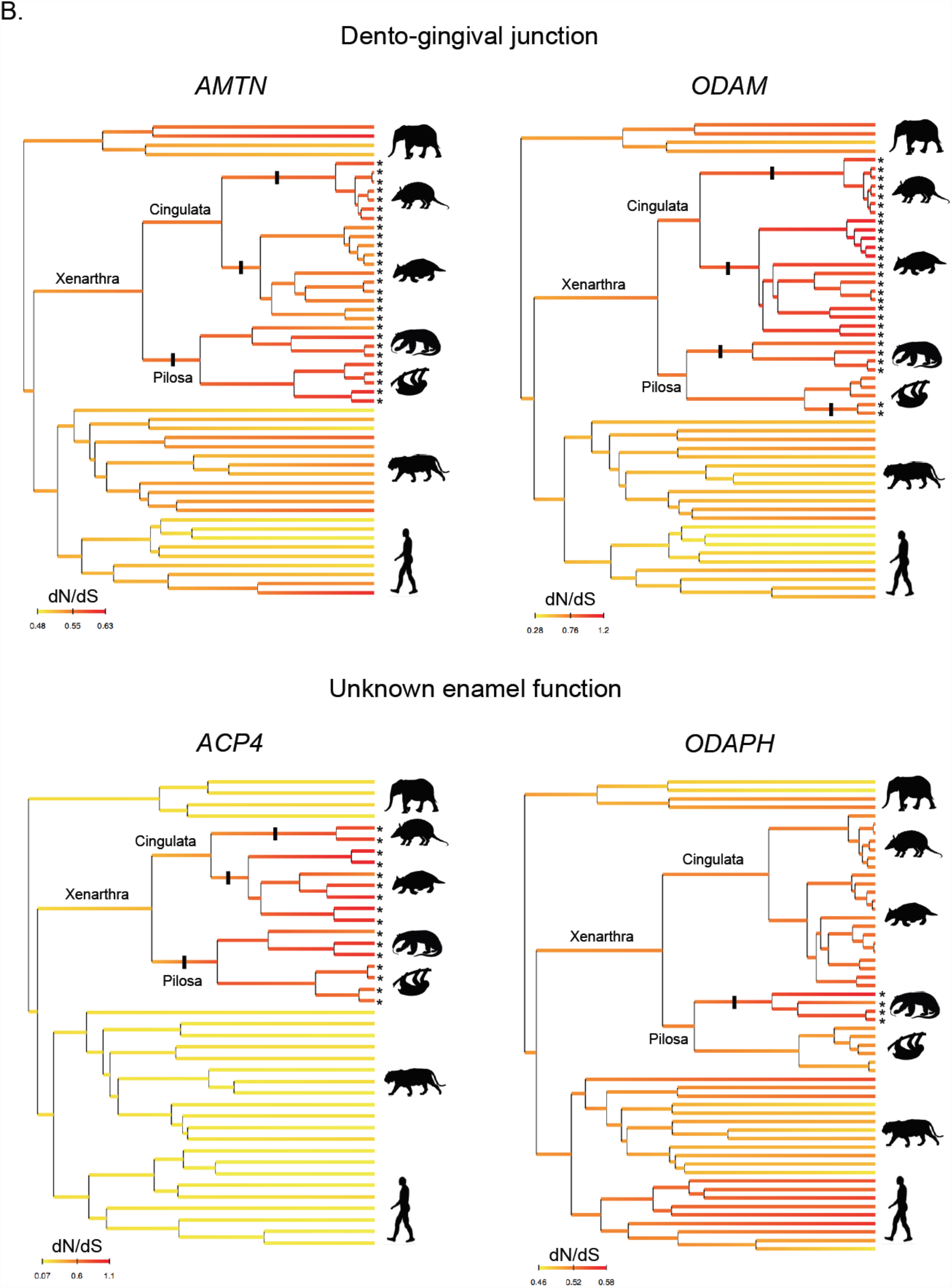
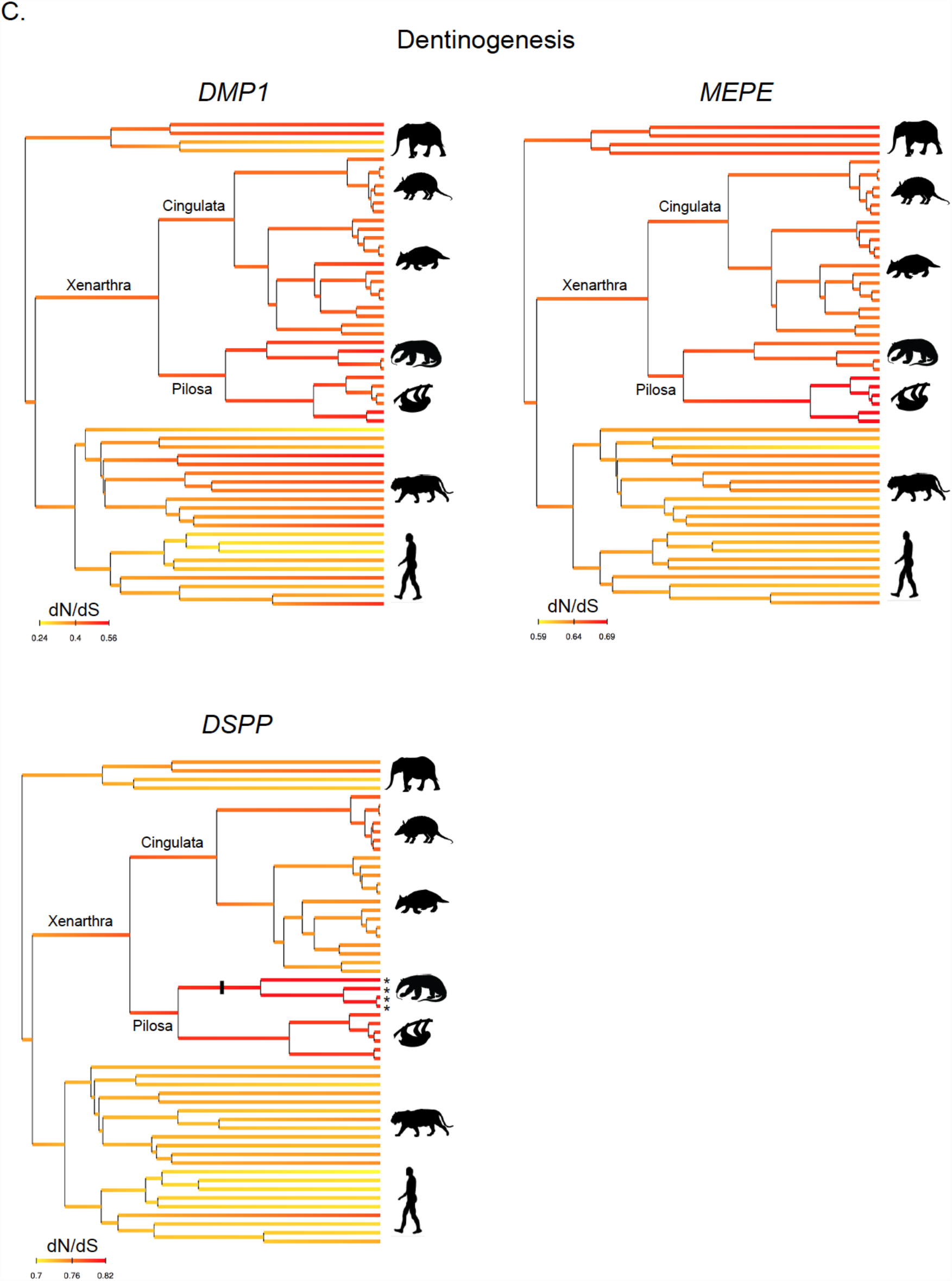
Bayesian reconstruction of dN/dS for 11 dental genes across the placental phylogeny with focus on xenarthrans (armadillos, anteaters, and sloths). A. Enamel matrix (*AMELX, AMBN, ENAM*) and enamel processing (*MMP20*) genes. B. Dento-gingival junction (*AMTN, ODAM*) and unknown enamel function (*ACP4, ODAPH*) genes. C. Dentinogenesis genes (*DMP1, MEPE, DSPP*). Asterisks at the tips of terminal branches identify non-functional sequences (pseudogenes) and bars on branches indicate shared inactivating mutations (SIMs). Detailed trees are provided as Supplementary Figures S25-S35. Silhouettes were obtained from PhyloPic (http://phylopic.org/).

The reconstructed patterns of relaxed selection were similar among genes encoding enamel matrix proteins (*AMELX, AMBN, ENAM*) (Figure 4A), with generally elevated dN/dS ratios trending towards an ω of 1. Consistent with the patterns of pseudogenization, *MMP20* shows elevated ω in pilosans, but distinctly lower estimates in armadillos (Figure 4A). Among the dento-gingival junction genes (*AMTN, ODAM*) (Figure 4B), *ODAM* shows a pattern similar to the enamel matrix proteins, whereas *AMTN* does not trend as strongly towards relaxed selection, despite evidence of pseudogenization in all xenarthrans. Among genes with unknown enamel functions, *ACP4* shows perhaps the greatest contrast between xenarthrans (relaxed selection) and the outgroup taxa (purifying selection), whereas *ODAPH* has more muted differences between the pseudogenes in vermilinguans and other branches (Figure 4B). For the dentinogenesis genes, the elevated pilosan *DSPP* branches contrast with other branches, whereas *DMP1* and *MEPE* seem to suggest minimal differences between xenarthran and other placental mammal branches (Figure 4C). Moreover, these analyses revealed that some transitional branches where gene inactivation was inferred based on SIMs have elevated dN/dS values. This was particularly evident for the Pilosa ancestral branch in which we identified SIMs in many different genes (*AMELX, ENAM, MMP20, AMTN*, and *ACP4*) (Figure 4A,B). Finally, these results helped pinpoint potential shifts in relaxed selection on branches predating the occurrence of SIMs such as the Cingulata ancestral branch in *AMTN, ODAM*, and *ACP4* in which the two main armadillo families (Dasypodidae and Chlamyphoridae) presented evidence of independent SIMs (Figure 4B).

While the Coevol results suggested potential shifts in selection on branches predating the occurrence of SIMs, we tested whether such inferences were supported statistically by likelihood ratio tests. We therefore performed dN/dS branch model analyses using codeml in PAML to estimate the selective pressure experienced by the 11 dental genes on branches that predate the timing of inactivating mutations in xenarthrans (Supplementary Tables S16-S26; Supplementary Figures S14-S24). These branch model analyses (Yang, 1998; Yang and Nielsen, 1998) estimate how natural selection is acting on a gene by comparing the ratio (ω) of nonsynonymous substitutions (dN) to synonymous substitutions (dS) accumulated in a gene on a given branch or set of branches on a phylogeny. ω < 1 suggests conservation of protein sequence (purifying selection) on average across the gene on that branch, ω > 1 is consistent with change in protein function (positive selection), and ω = 1 is associated with relaxed selection, which is the pattern expected for pseudogenes. However, on a phylogenetic branch that has a mixed history, such as purifying selection followed by relaxed selection, ω should be intermediate between the average strength of selection and one (Meredith et al., 2009). As such, we estimated the average background ω, and tested whether key branches were statistically elevated or lowered compared to this background selection pattern and an ω of 1.

We first tested whether ω was ever elevated on branches that predated the earliest SIMs and found several such examples. For instance, *AMELX* possesses a SIM for *Cabassous* + *Tolypeutes*, with the sister taxon *Priodontes* only possessing unique mutations. However, the branch immediately ancestral to these armadillos (stem Tolypeutinae) has an elevated ω (2.29, versus background of 0.33 [p = 0.035]), suggesting relaxed selection on *AMELX* began on this branch. Similarly, *ENAM* has a SIM for *Cabassous* + *Tolypeutes*, but not for *Priodontes*, but again model comparisons suggest relaxed selection began on the stem Tolypeutinae branch (ω = 1.21, versus background of 0.52 [p = 0.022]).

Given such examples, we tested whether elevated ω estimates could be found on the two xenarthran branches that predate any SIMs: stem Xenarthra and stem Cingulata. For the stem xenarthran branch, we found a single example where a gene had ω statistically distinguishable from the background (*ACP4* = 0.57, versus background of 0.18), although this was only marginally significant (p = 0.049). By contrast, five of the 11 genes had signals consistent with purifying selection (*AMELX* [ω = 0.0001], *DMP1* [ω = 0.37], *ENAM* [ω = 0.48], *MMP20* [ω = 0.2], *ODAPH* [ω = 0.14]), being statistically distinguishable from 1 but not from the background. *ODAPH* had an ω so low that it was statistically distinguishable from both 1 and the background (ω = 0.48). For the stem Cingulata branch, 10 of the 11 genes showed no evidence of elevated ω, with five instead being consistent with purifying selection (*ACP4* [ω = 0.1], *AMBN* [ω = 0.17], *DMP1* [ω = 0.34], *MMP20* [ω = 0.12], *ODAPH* [ω = 0.32]). For *AMBN*, ω was so low that it was distinct from the background (ω = 0.41). The one example of an elevated ω was *DSPP* (ω = 1.46), though we found no evidence of inactivation in this dentin-specific gene among the dentin-retaining armadillos.

## Discussion

### Reconstructing the dental regression history of xenarthrans

Our results suggest that certain major regressive events in the dental history of xenarthrans took place independently and in a gradual, stepwise fashion. Four lines of evidence suggest that this regression occurred in parallel in the three main xenarthran lineages rather than in their last common ancestor. First, among the nine genes that would ultimately become pseudogenes in at least some xenarthrans, no unambiguous shared inactivating mutations (SIMs) were inferred for the entire xenarthran clade. Second, no SIMs were found between the two major armadillo clades despite the last common ancestor of cingulates dating to roughly 23 million years after the origin of Xenarthra (Gibb et al., 2016). Third, while we found evidence of elevated dN/dS for a single gene (*ACP4*) on the stem xenarthran branch, this estimate was only marginally significant and the descendant Cingulata branch showed purifying selection on this gene. By contrast, we found five genes showing statistically significant purifying selection on this branch, three of which are enamel-specific genes with widespread patterns of pseudogenization in vertebrates with dental regression (*AMELX, ENAM, MMP20*) (Meredith et al., 2009, 2011a, 2013, 2014; Choo et al., 2016). Fourth, the same can be said for the stem armadillo branch, on which five genes were also statistically consistent with purifying selection, with only the dentin-specific *DSPP* having a statistically elevated ω, despite never being inactivated in any armadillos.

These four lines of evidence predict that the last common ancestor (LCA) of xenarthrans, the stem-cingulate lineage, and the LCA of Cingulata all had teeth covered with enamel, with descendant lineages subsequently deriving thin enamel (dasypodid armadillos), losing enamel (chlamyphorid armadillos, sloths) or losing teeth entirely (anteaters). However, this does not negate the possibility, nor indeed the likelihood, that some degree of tooth simplification had already occurred by the last common ancestor of xenarthrans and/or armadillos. Notably, the two earliest armadillo fossils that preserve teeth, which are roughly coeval with molecular estimates of the LCA of cingulates (Gibb et al., 2016), have simple, peg-like teeth with thin enamel, although they differ in whether the enamel wore easily (*Utaetus buccatus* [42–39 Mya]) or not (*Astegotherium dichotomus* [45 mya]). Notably, the former is a putative chlamyphorid relative, and the latter is considered a relative of dasypodids (Ciancio et al., 2014; Simpson, 1932). Their enamel is reminiscent of phenotypes that can arise from amelogenesis imperfecta (AI), a condition that can lead to the formation of thin, soft and/or brittle enamel that wears away with time, which can be caused by the inactivation a number of the numerous enamel-associated genes (Smith et al., 2017). It remains possible that non-coding mutations leading to dental regression could have accumulated prior to the mutations more typically characteristic of pseudogenes, at least by the origin of cingulates, a potentially fruitful avenue for future research.

After the origin of cingulates, our data suggest that *AMTN, ODAM* and *ACP4* were independently inactivated in stem chlamyphorid and stem dasypodid armadillos by at least 37 Mya and 12 Mya (Figures 1 and 3), respectively (Gibb et al., 2016). The proteins produced by the former two genes, AMTN and ODAM, are both expressed in enamel-producing ameloblasts (Fouillen et al., 2017), but their link to enamel integrity is tenuous. Though the deletion of exons 3-6 in *AMTN* has been linked to AI in humans (Smith et al., 2016), mouse knockouts display a minimally-affected dental phenotype (Nakayama et al., 2015). Specifically, mandibular incisors have chalky enamel, which begins to chip away after 13 weeks of age, but maxillary incisors and molars show no significant alterations. *ODAM* knockout mice do not seem to have affected enamel whatsoever (Wazen et al., 2015), and this gene has not been implicated in enamel malformations in humans (Smith et al., 2017). Both proteins, however, continue to be expressed throughout adulthood in the junctional epithelium (Ganss and Abbarin, 2014). These proteins appear to congregate into an extracellular matrix to maintain a tight seal between the gingiva and teeth (Fouillen et al., 2017), presumably to protect the teeth from microbial exposure (Lee et al., 2015). Notably, *ODAM* knockout mice show a decreased ability to heal the junctional epithelium after damage (Wazen et al., 2015), lending credence to this hypothesis. *ACP4’*s function is less well-known, but it is expressed during amelogenesis (Seyman et al., 2016) and disabling mutations in this gene leads to hypoplastic AI, leading to thin enamel (Kim et al., 2022; Liang et al., 2022; Seyman et al., 2016; Smith et al., 2017). Accordingly, the losses of both *AMTN, ODAM* and *ACP4* in stem chlamyphorids and stem dasypodids, respectively, point to a weakening of the gingiva-tooth association and a thinning of enamel during the origins of these subclades.

After the origin of chlamyphorids (37 Mya), we inferred that further pseudogenizations of *AMBN, AMELX*, and *ENAM* took place, apparently in parallel (Figure 1). Disabling any of these three genes, all of which are expressed in ameloblasts, is associated with AI (Lagerström et al., 1991; Poulter et al., 2014; Rajpar, 2001; Seymen et al., 2016), leaving little doubt that enamel regression and eventually loss occurred during the diversification of this clade. Curiously, *MMP20* appears universally intact in chlamyphorids but shows evidence of elevated dN/dS estimates, possibly indicating a change in function for this gene that is otherwise strongly linked to enamel development (Meredith et al., 2011a, 2013, 2014; Smith et al., 2017).

By contrast, the comparably late and minimal degree of pseudogenization in dasypodids may explain the retention of vestigial enamel on the milk teeth and thin, easily worn enamel on the permanent teeth of dasypodids (Ciancio et al., 2021; Martin, 1916; Spurgin, 1904). After the stem loss of *AMTN, ODAM* and *ACP4, ENAM* is the only gene with clear evidence of pseudogenization in multiple dasypodids, implying a delay in the historical timing of dental degeneration compared to chlamyphorids. The pattern of gene inactivation in dasypodids is notable in that *ENAM* is inactivated in several species and *Dasypus novemcinctus* has pseudogenic *ENAM, AMBN* and *ACP4*, yet thin regressive enamel is still present in these species (Ciancio et al., 2021; Martin, 1916; Spurgin, 1904). This suggests that inactivation of these genes individually or in concert is insufficient for complete enamel loss. Notably, *AMELX* appears intact in all dasypodids, and was under purifying selection on the stem dasypodid branch, yet is pseudogenic in all pilosans and chlamyphorids sampled. This implies that *AMELX* may be a strong indicator for the timing of enamel loss in xenarthrans, a possibility made more plausible by its critical function (see below). If correct, enamel loss appears to have occurred up to five times independently within Chlamyphoridae based on SIMs (Figure 1).

In Pilosa, our results suggest that the first eight million years of their history (Gibb et al., 2016) resulted in a relatively rapid regression of their dentition. Specifically, we inferred that *AMTN, AMELX, ENAM, MMP20* and *ACP4* all became pseudogenes prior to the last common ancestor of pilosans (Figures 1 and 2). Enamel-forming ameloblasts express the enamel matrix proteins (EMPs) AMELX and ENAM, both of which are processed into their mature peptide forms by the metalloproteinase MMP20 (Smith et al., 2017). Building off an earlier study of *ENAM* (Meredith et al., 2009), this strongly suggests that enamel was completely lost by the earliest pilosans, particularly given that AMELX makes up to 90% of the EMP composition and MMP20 is critical for processing the EMPs AMELX, AMBN and ENAM (Smith et al., 2017). The earliest tooth-bearing fossil pilosan is a stem sloth (*Pseudoglyptodon*) dating to the early Oligocene (32 Mya) (McKenna et al., 2006), well after the estimate for the earliest pilosans (58 Mya) (Gibb et al., 2016). As predicted, *Pseudoglyptodon* lacks enamel, but our results predict the future discovery of enamelless stem and/or crown pilosans in the Paleocene.

Our results further suggest that stem vermilinguans inherited this enamelless condition, and then continued dental regression to the point of complete tooth loss. In addition to all modern anteaters being edentulous, we found a putative shared mutation in *ODAPH* that suggests complete tooth loss occurred by the origin of Vermilingua (38 Mya) (Gibb et al., 2016). Although *ODAPH* is considered to be important in enamel formation (Parry et al., 2012; Prasad et al., 2016), a comparative study of this gene in placental mammals suggested that it is uniquely inactivated in the edentulous baleen whales and pangolins (Springer et al., 2016) Our results support the hypothesis that *ODAPH* has a critical function outside of enamel formation. Perhaps more convincing is the evidence suggesting pseudogenization of the dentin-matrix protein *DSPP* in a stem vermilinguan (Figure 2). Inactivation of *DSPP* leads to dentinogenesis imperfecta in humans (Xiao et al., 2001) and dentin defects in knockout mice (Sreenath et al., 2003), increasing the likelihood that tooth loss occurred prior to the origin of this clade. The earliest known anteater, the edentulous *Protamandua* (Gaudin and Branham, 1998), dates back to the Santacrucian (17.5–16.3 Mya), but our results predict toothless stem vermilinguans likely dating to the Eocene, between 58 and 38 Mya (Gibb et al., 2016).

### Broader implications for regressive evolution

Our results reveal a few noteworthy patterns that may inform the study of regressive evolution in other systems. First, these data provide genomic evidence that trait loss can take place in a stepwise manner. Fossil evidence implies that both turtles and birds lost their teeth gradually: turtles first lost marginal dentition, followed by palatal teeth (Li et al., 2008), and birds lost premaxillary teeth prior to becoming completely edentulous (Meredith et al., 2014). Xenarthran dental genes point to a third path towards this phenotype, with discrete events of enamel loss followed by tooth loss in the evolution of anteaters, based on shared and distinct pseudogenization signals in vermilinguans and sloths. This dental regression scenario was also recently inferred in a study on the evolution of baleen whales (Mysticeti) based on a similar comparison of dental genes (Randall et al., 2022).

Second, our results suggest that regressive evolution can vary broadly in timespan and pattern. Although we inferred that enamel loss occurred in stem pilosans within an eight million year window, dasypodid dentition, by contrast, appears to be the product of a much lengthier period of regression. The earliest putative crown cingulate fossils have simplified teeth with thin enamel and date back to 45 Mya (Ciancio et al., 2014). Assuming that these taxa represent the ancestral condition for crown armadillos, then the loss of *ODAM, AMTN* and *ACP4* in a stem dasypodid, inactivation of *ENAM* in several crown dasypodids, and individual examples of *MMP20* and *AMBN* pseudogenization, imply a protracted and evidently episodic, pattern of regression in this lineage.

Finally, these data provide insights into how the regression of traits, such as the weakening or wholesale loss of enamel, may constrain the evolutionary trajectories lineages can take. The simplification and loss of teeth is a relatively common phenomenon in mammals, with most such species having diets characterized as being myrmecophagous (ants and/or termites), vermivorous (soft-bodied worms) or nectarivorous (Charles et al., 2013; Davit-Béal et al., 2009; Freeman, 1995; Rosenberg and Richardson, 1995). Presumably the simplification or loss of teeth in such taxa is due to the softness of their prey (vermivores) or reliance on the tongue for food acquisition and minimized need for mastication (myrmecophagy, nectarivory). Notably, vermilinguan xenarthrans are myrmecophagous, and most armadillos at least partially, and at most extensively, consume social insects (Nowak, 1999), so possessing simplified teeth or being edentulous likely reinforces this diet. Reconstructions of chitinase genes in the earliest pilosans suggest that they were likely highly insectivorous (Emerling et al., 2018), indicating myrmecophagy is a plausible explanation for their early dental regression. However, after deriving enamelless teeth in stem pilosans, sloths predominantly became herbivores (Nowak, 1999; Saarinen and Karme, 2017). In contrast to the simplified, enamelless teeth present in sloths, other extant herbivorous mammals have enamel-capped teeth that tend towards an increased complexity of tooth cusps (Ungar, 2010). Yet, whereas anteaters continued the trend towards dental simplification with complete tooth loss, natural selection likely strongly favored the retention of teeth in sloths, as evidenced by the signal of purifying selection found in *ODAPH* on the stem sloth branch, with elevated dN/dS values on the dentin-associated genes *DMP1* and *MEPE* (Figure 4; Supplementary Figures S18, S21, S24) potentially pointing to positive selection resulting in functional changes to sloth dentin.

In summary, our results show how pseudogenes can provide insights into the deep evolutionary history of a clade, and point to the divergent paths that regressive evolution can take. Understanding of this system would be enhanced by analyzing non-coding elements and functional data, given that mutations outside of the protein-coding regions of these genes may pre-date frameshift indels, premature stop codons, and similar inactivating mutations. Furthermore, as new tooth-specific genes are discovered, they may provide further resolution to our understanding of this question.

## Acknowledgements

We would like to thank Anaïs Tibi and Delphine Sérol for their contribution to this study. We also thank Sérgio Ferreira-Cardoso and Lionel Hautier for helpful discussions. Jeremy Johnson (Broad Institute, Cambridge, MA, USA) kindly provided early access to xenarthran genome assemblies. This work would not have been possible without the help of the following individuals and institutions in accessing xenarthran tissue samples: Mariella Superina, Jim Loughry, Agustín Jiménez, Rodolfo Rearte, Guido Valverde, and Guillermo Pérez-Jimeno; Philippe Gaucher, Roxane Schaub, Gérard Lievin, Erika Taube, Philippe Cerdan, Michel Blanc, Maël Dewynter, Sébastien Barrioz, Rodolphe Paowé, Jean-François Mauffrey, Eric Hansen, François Ouhoud-Renoux, and Jean-Christophe Vié (French Guiana); Baptiste Chenet and David Gomis (Zoo de Lunaret, Montpellier, France); John Trupkiewich (Philadelphia Zoo, USA); Daniel Hernández (Facultad de Ciencias, Universidad de la República, Montevideo, Uruguay); Sergio Vizcaíno (Museo de La Plata, La Plata, Argentina); Ross MacPhee (American Museum of Natural History, New York, USA); Jonathan Dunnum and Joseph Cook (Museum of Southwestern Biology, Albuquerque, USA); Jake Esselstyn, Donna Dittman, and Mark Hafner (Louisiana State University Museum of Natural Science, Baton Rouge, USA); Darrin Lunde (National Museum of Natural History, Washington, USA); Jim Patton (Museum of Vertebrate Zoology, Berkeley, USA); Gerhard Haszprunar and Michael Hiermeier (The Bavarian State Collection of Zoology, Munich, Germany); Benoit de Thoisy (Kwata NGO and Institut Pasteur de la Guyane, Cayenne, French Guiana); Géraldine Véron and Violaine Nicolas (Museum National d’Histoire Naturelle, Paris, France); and François Catzeflis (Institut des Sciences de l’Evolution, Montpellier, France). We thank Didier Casane, Juan Opazo, Régis Debruyne, and Nicolas Pollet for helpful comments during the Peer Community in Genomics recommendation process. Sanger sequencing data were produced through technical facilities of the SeqGen platform of the Labex CeMEB (Centre Méditerranéen Environnement Biodiversité). Phylogenetic and statistical analyses benefited from the Montpellier Bioinformatics Biodiversity platform (MBB). This is contribution ISEM-2023-XXX of the Institut des Sciences de l’Evolution.

## Data, scripts, code, and supplementary information availability

Data are available online: https://doi.org/10.5281/zenodo.8005757

Scripts and code are available online: https://doi.org/10.5281/zenodo.8005757

Supplementary information is available online: https://doi.org/10.5281/zenodo.8005757

